# Unraveling *Streptococcus pyogenes* Carriage: Genomic and Transcriptomic Insights from Acute and Post-Treatment Phases

**DOI:** 10.1101/2025.10.22.683982

**Authors:** Nicholas Faiola, Jason Woods, May Xu, Kevin Quirk, Christopher Thomas, Eliza Klos, Jordan Thesier, Mitchell Waldran, Mark Mainschein-Cline, Michael J. Federle, Gregory P. Demuri, Ellen R. Wald, Laura Cook

**Affiliations:** Department of Biological Sciences, Binghamton Biofilm Research Center, Binghamton University, Binghamton, New York, USA; Core for Research Informatics, Research Resources Center, University of Illinois at Chicago, Chicago, Illinois, USA; Department of Medicinal Chemistry and Pharmacognosy, University of Illinois at Chicago, Chicago, Illinois, USA; Department of Pediatrics, Division of Infectious Diseases, University of Wisconsin School of Medicine and Public Health, Madison, WI, USA

## Abstract

The carrier state of *Streptococcus pyogenes* (Group A Streptococcus, GAS), is defined by the presence of the organism without symptoms or an immune response. Carriage poses challenges for clinical management and contributes to transmission in the community. Prior studies suggest genetic and phenotypic differences between isolates from symptomatic (acute) infections and those from carriers, yet comprehensive genomic and transcriptomic analyses of naturally acquired human GAS carriage have been lacking. In this study, we collected longitudinal samples from individuals before and after antibiotic treatment for acute pharyngitis, performing whole genome sequencing and RNA-seq on selected isolates. This is the first report of transcriptional profiling of GAS directly from human derived oropharyngeal swabs. Genomic analysis revealed no consistent carrier-specific genotypes, and *in vitro* assays showed no major differences in biofilm formation or antibiotic susceptibility between carrier and acute isolates. Transcriptional profiling from oropharyngeal swabs identified distinct gene expression patterns when comparing the acute infection and post-treatment carriage phases, although early acute expression profiles did not predict treatment failure. These findings suggest that GAS carriage is likely influenced by conserved bacterial traits as well as host factors, advancing our understanding of GAS persistence and transmission.

**Summary:** Longitudinal samples were collected from individuals before and after treatment for acute Group A Streptococcal pharyngitis. Whole genome sequencing and RNA-seq was performed on selected isolates. This is the first report of transcriptional profiling of GAS directly from human oropharyngeal swabs. Distinct gene expression patterns were observed when comparing the acute infection and post-treatment carriage.

## Introduction

Many species of bacteria that colonize the human upper respiratory tract maintain dichotomous lifestyles, capable of both commensal and pathogenic relationships with the host. Species such as *Staphylococcus aureus, Streptococcus pneumoniae* and *Streptococcus pyogenes* (Group A Streptococcus, GAS) cause considerable clinical and economic burdens due to frequent infections; yet these species are carried asymptomatically in the upper respiratory tract of the human population at high rates. GAS is the most common cause of bacterial pharyngitis in children and adults and causes both suppurative (acute otitis media, acute sinusitis, mastoiditis, musculoskeletal and systemic infections) and non-suppurative (post-streptococcal glomerulonephritis and rheumatic fever) sequelae. Every year, over 600 million new cases of GAS pharyngitis occur worldwide (1). Rates of symptomatic GAS pharyngitis vary between 15-30% for children and 4-10% in adults in the developed world with even higher rates in developing countries (1–6). In the last decade, there has been a documented re-emergence of both superficial and invasive *S. pyogenes* infections worldwide (7–9).

Most superficial GAS infections are cleared spontaneously or after appropriate antibiotic administration; however, in many children, GAS becomes part of the normal pharyngeal microbiota, leading to a GAS carrier state. The carrier state of GAS is defined as the presence of the organism in the pharynx without evidence of inflammation or increased serological response to extracellular antigens. Pharyngeal GAS carriage rates in asymptomatic school-aged children vary based on location but are most often estimated to be between 8-20% (4, 10–13). It has been proposed that the true streptococcal carrier state can be most accurately determined retrospectively as antimicrobial treatment failure following an acute infection (14), and several studies have used treatment failure to define GAS carriers (15–17).

The carrier state is of particular significance to public health because carriers may serve as a reservoir for the spread of GAS to others in the population and could harbor accessory genetic information that could be acquired through horizontal transfer to co-colonizing strains and species. GAS carriage has been associated with community outbreaks of pharyngitis (18) and fatal invasive disease in close contacts (19), and treatment failure has been implicated in development of post-infectious sequelae such as acute rheumatic fever (20). In addition, the carrier state is a confounder for clinicians. Patients with viral pharyngitis who are also carriers may test positive for GAS, leading to antibiotic administration and an estimated 1.5 million unnecessary antibiotic courses annually in the United States (14). It is also likely that the inflammation in the upper airway associated with symptoms of acute bacterial or viral pharyngitis is associated with increased transmission of GAS by carriers.

Previous studies have found that some cultured GAS isolates recovered from asymptomatic individuals contained genetic polymorphisms that, when integrated into a virulent isolate, caused a carrier-like phenotype in animal models (21–23). These findings point to asymptomatic carriage being the result of genetic mutations, potentially arising during virulent episodes, that effectively diminish virulence but maintain survival in the host, at least temporarily. This contradicts conventional thinking in relation to other pathobionts like *S. aureus* and *Haemophilus influenzae*, in which the carried bacterium is considered to be the evolutionary ideal due to survival advantage. It is therefore important to thoroughly test these potentially contradictory hypotheses using naturally occurring and empirically determined carrier isolates from human patients.

To date, no studies have examined clinical isolates obtained longitudinally from an acute pharyngitis event through to the carrier state at genomic and transcriptomic levels. Transcriptional analysis of previously obtained carriage isolates has not previously been possible as these strains have all been cultured, changing transcriptional patterns. Swabs directly from patients either during or after an acute infection have not been examined via RNA-seq due to technical challenges in obtaining and processing these samples. An important study conducted in a non-human primate macaque model of long-term GAS upper respiratory tract infection delineated gene expression in GAS during the course of >80 days (24) but no transcriptional analysis of natural human acute and carrier samples has been undertaken.

Here we describe collection of longitudinal samples from human subjects treated for acute streptococcal pharyngitis both before and after antibiotic treatment. We performed whole genome sequencing and transcriptional profiling of selected isolates, comparing patient swabs that were cleared of GAS by treatment (acute samples), with those that remained colonized after treatment (carriers).

Whole genome sequencing did not identify conserved carrier-specific genotypes and *in vitro* analysis of biofilm formation and antibiotic resistance profiles showed no major differences between acute and carrier isolates. Transcriptional profiles differ between GAS present in the oropharynx during the acute and post-antibiotic carriage stage of infection. Transcriptomics comparing acute and carrier isolates during the acute phase of infection did not identify expression signatures predicting future treatment failure. Together these results provide insight into strain-specific carriage potential and indicate a larger role for the host environment and/or host genetics in determining GAS carriage.

## Results

### Collection of longitudinal patient samples

A flow chart of the sample collection process is shown in Figure 1. Samples used in this study were collected from children recruited from four pediatric clinics in Madison, WI between 2014-2023.

**Figure 1.**
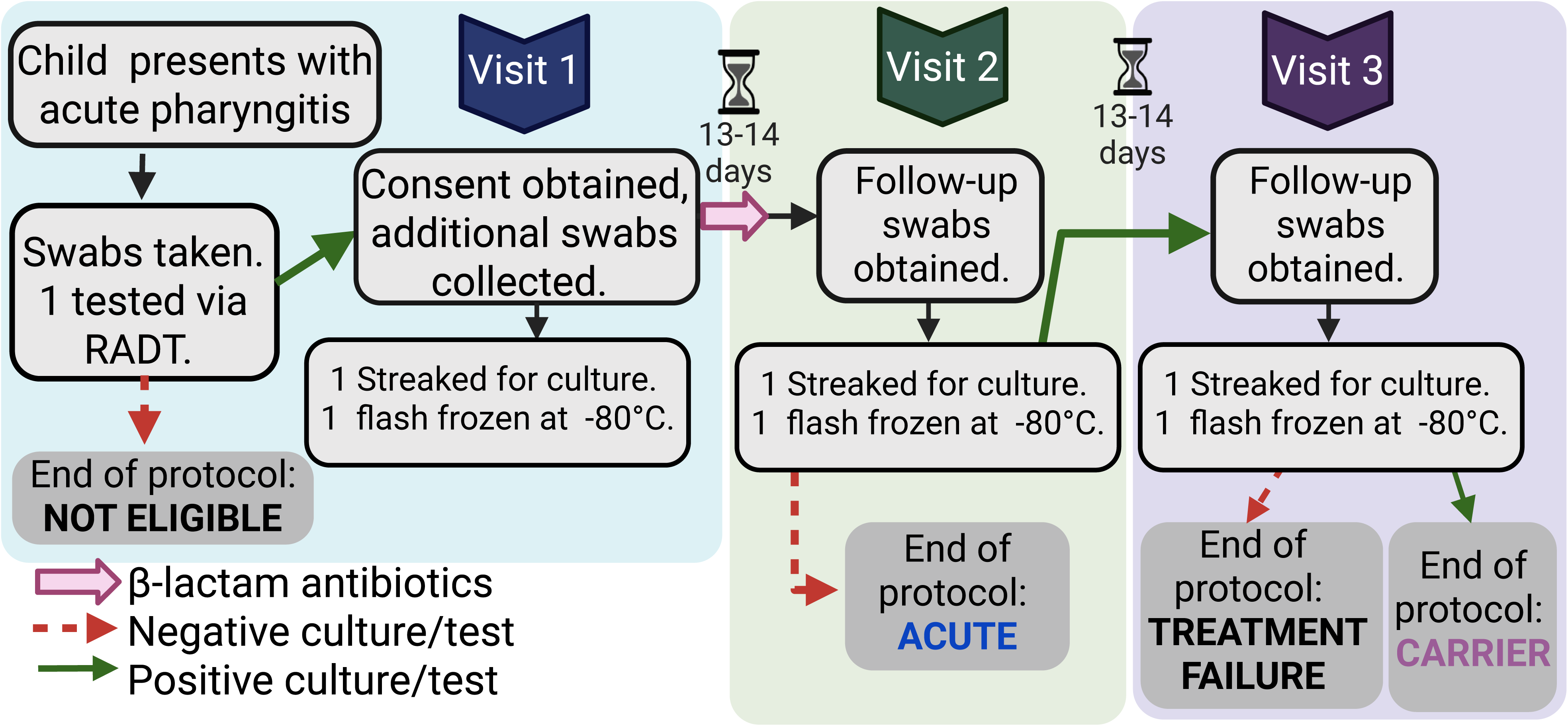
Sample collection diagram outlining the patient swab collection protocol. Flowchart of study participants and acute and carrier swab sample collection. Eligible participants are between 5-15 years of age, not allergic to β-lactam antibiotics, and positive for GAS using a rapid antigen detection test (RADT). Swabs are collected during the initial visit and after a course of β-lactam antibiotic treatment. Patients with a positive throat culture for GAS 20-24 days after the initial visit are considered GAS carriers and isolates identified as carriage isolates (Car). Patients who are negative on GAS throat culture at this point are assumed to have had an acute infection and termed Acute (Act) in future experiments. Red arrows indicate a negative test and green arrows a positive test. Figure created in Biorender.

Eligibility criteria included age between 5 and 15 years of age, a diagnosis of acute pharyngitis, and having not been treated with antibiotics in the last 30 days. Children were excluded from this study if they had an allergy to β-lactam antimicrobials or if they were treated with an antibiotic other than a β-lactam.

Children presented to the clinics with acute symptoms of sore throat (visit 1). A swab was obtained from the posterior pharynx (tonsillar pillars and posterior pharyngeal wall) of the patient. A rapid antigen detection test (RADT) was used for initial diagnosis. If the RADT was positive, the child and family were informed by the research staff that the child was eligible for the study and if the family and patient agreed to participate, signed informed consent was obtained and two additional throat swabs were obtained. One was stored immediately at -80 °C while another was forwarded to the laboratory to culture for GAS using standard methods and the cultured isolate preserved. Treatment was then undertaken with an appropriate dose of either amoxicillin or penicillin V for up to 10 days at the discretion of the provider. Following the course of antibiotics, patients returned for a follow-up in the clinic at day 14 (visit 2). If the child was asymptomatic and lacked inflammation of the pharynx or tonsils, a double swab was used to sample the posterior pharynx. If the follow-up swab was negative for GAS on culture, the patient was deemed to have presented with an acute infection cleared by antibiotics. If the follow-up swab was positive, the second swab (potential-carrier) was stored at -80 °C. An additional set of swabs was obtained approximately two weeks later (visit 3, approximately 4 weeks from initial presentation). If the swab was still positive for GAS by culture, the subject was deemed a carrier and the swab was again stored at -80 °C. All cultured isolates from carriers were tested by PCR to demonstrate concurrent *emm* types between the first and last swabs. The strains isolated and examined here are described in Table 1. Each isolate was marked as acute or carrier and carrier isolates were marked as being collected on the initial visit (-1) or the third and final return visit ∼4 weeks later (-3).

**Table 1:**
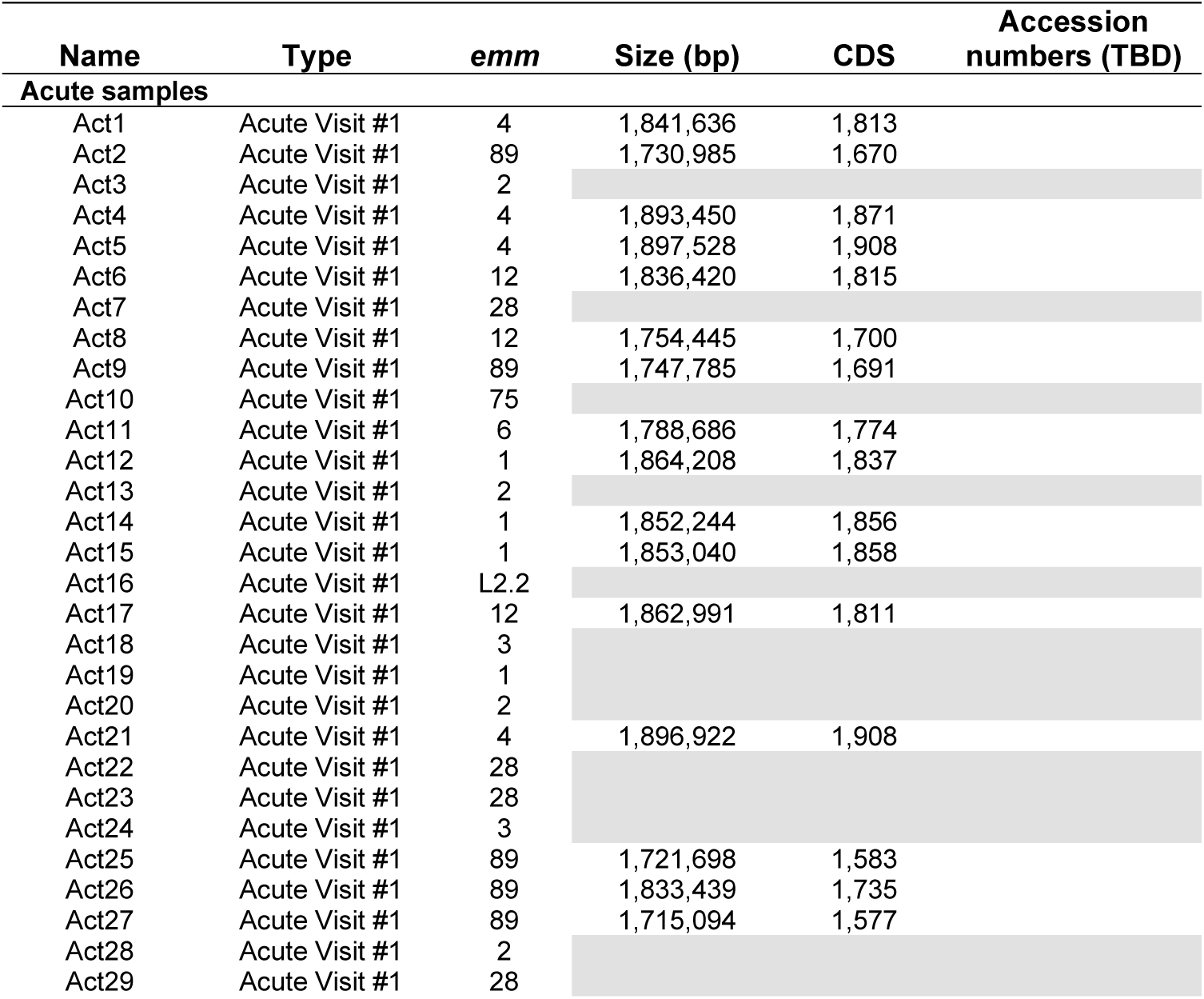
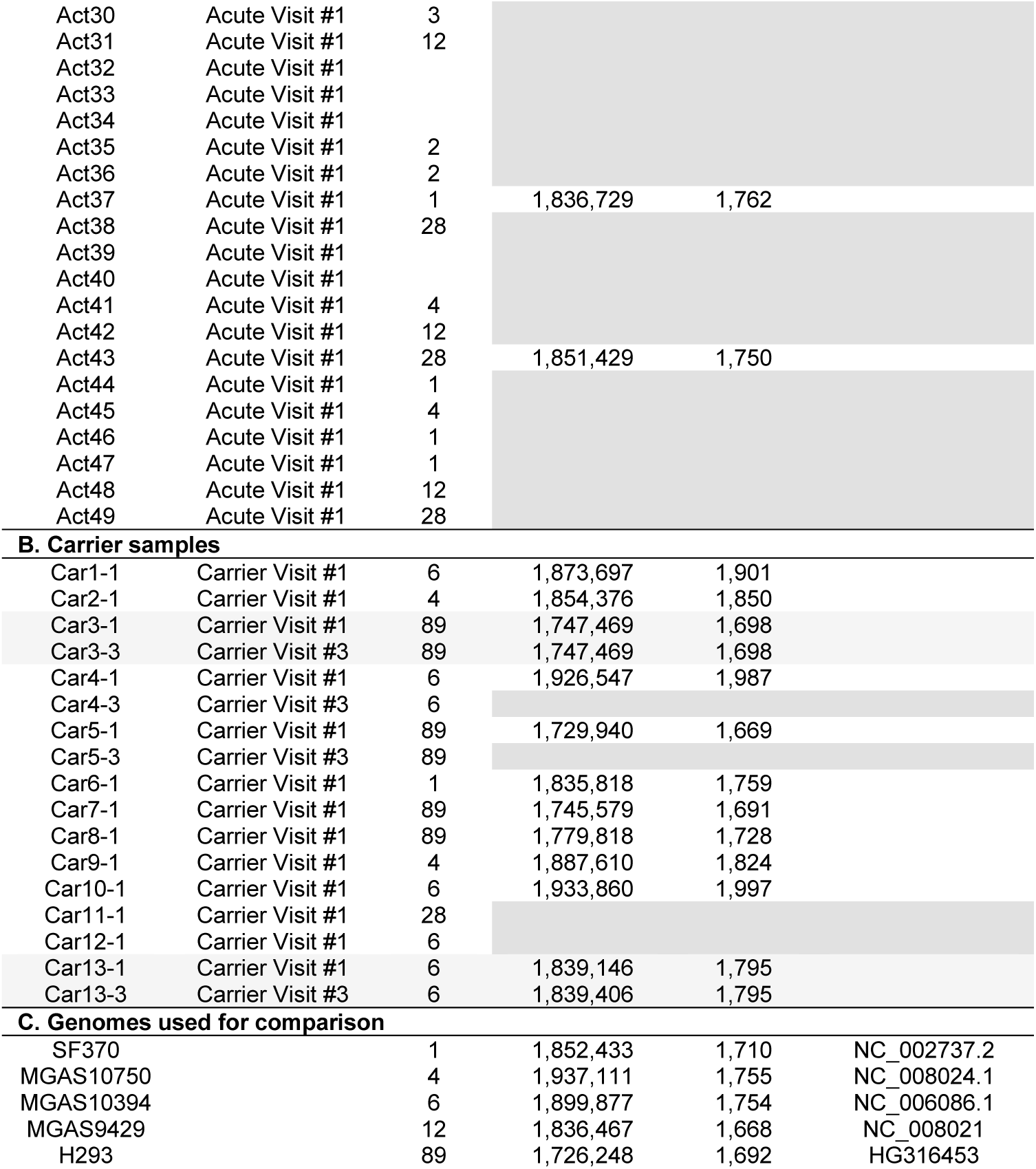
*Streptococcus pyogenes* isolates used in this study.

### Antibiotic resistance profiles

The minimum inhibitory concentration of four antibiotics was examined for 40 acute and 13 carrier strains using Liofilchem antibiotic test strips on blood agar plates (Table 2). All strains were identified as susceptible to penicillin and amoxicillin, the antibiotics used clinically for treatment of patients in this study, indicating that antibiotic resistance did not arise in the carrier strains even in those that remained following a course of antibiotics. Three of the collected isolates showed resistance to erythromycin, Car2 (*emm4*), Car10 (*emm6)*, and Act46 (*emm1*). One strain, Act39, was resistant to clindamycin.

**Table 2:** Antibiotic resistance profiles of clinical isolates.

| Antibiotic | R (%) | S (%) | MIC range |
| --- | --- | --- | --- |
| **β-lactam** |  |  |  |
| Penicillin | 0 | 100 | S = 0.003-0.047 |
| Amoxicillin | 0 | 100 | S = <0.016-0.047 |
| **Macrolide** |  |  |  |
| Erythromycin | n=3<br>(5.3%) | n=54<br>(94.7%) | S = 0.016-0.25<br>R>0.25 |
| **Lincosamide** |  |  |  |
| Clindamycin | n=1<br>(1.8%) | n=56<br>(98.2%) | S = 0.016-0.5<br>R>0.5 |

Acute n = 40, Carrier n = 17, 13 first visit, 4 return visit

Overall, very low rates of antibiotic resistance were observed in the collected isolates as expected and in this study all patients were treated with a beta-lactam.

### *emm* type distribution

57 cultured strains (44 acute patient strains and 13 carrier patients, 4 of which had initial and return visit cultures available and 9 of which had only initial visit cultures) were assessed for *emm* type using PCR and Sanger sequencing as described by the CDC (25). All carrier return isolates had identical *emm* sequences to the initial visit isolates so only one sample was included in *emm* type analysis to not skew the results. The distribution of *emm* types in these isolates are shown in Figure 2a. In this small cohort of samples, acute isolates had a wider range of *emm* types and a more even spread than carrier isolates. The percent of carrier isolates with *emm* type 6 was significantly higher than expected from the acute isolate data and the majority of the carrier isolates (9 of 13) were either *emm* type 6 or 89.

**Figure 2.**
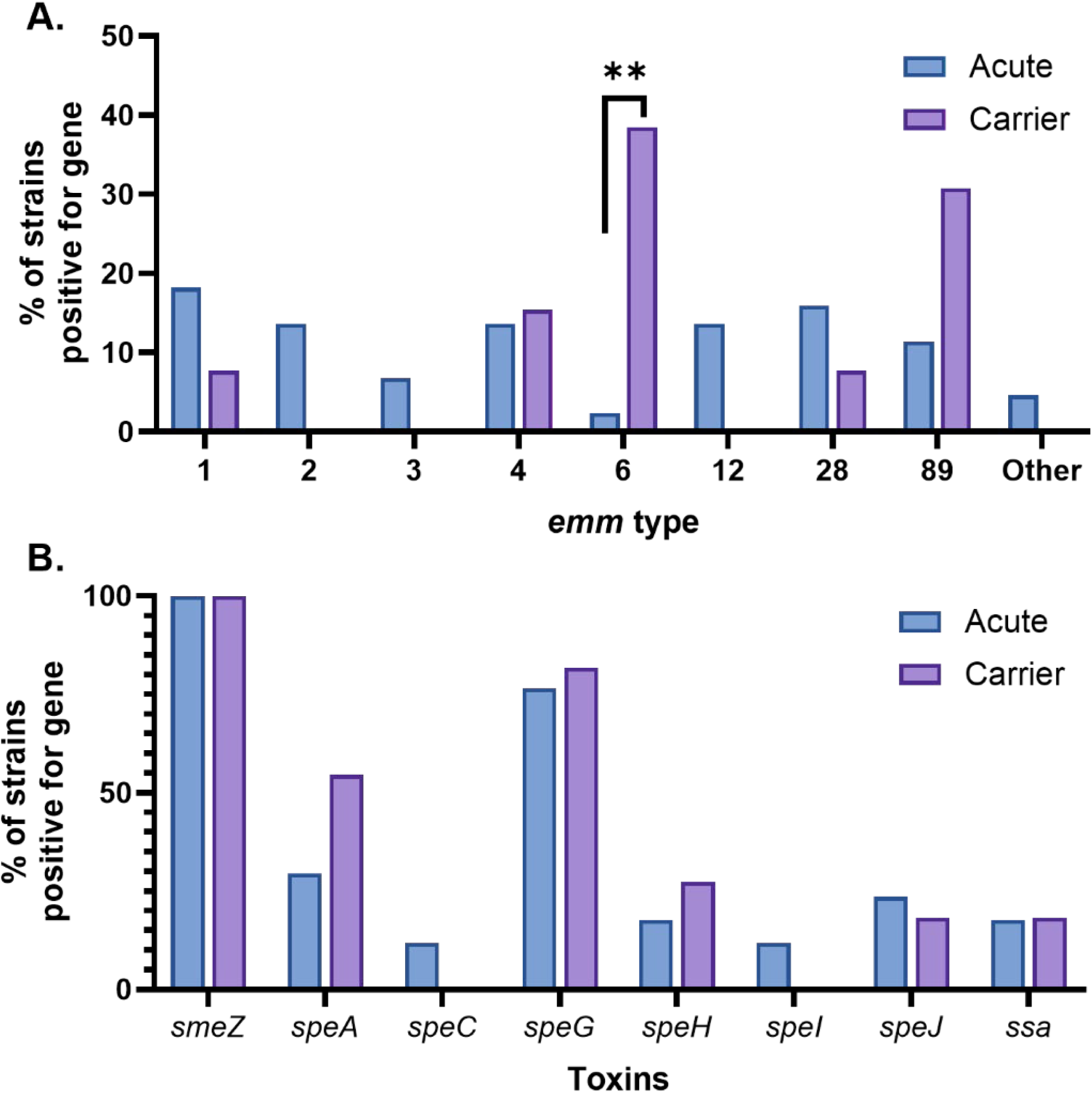
emm type and toxin profiles of acute and carrier GAS strains. (A) *emm* type distribution in acute (blue) and carrier (purple) isolates. Fisher’s exact test was used to determine statistical significance. ** p value < 0.01. (B) Distribution of toxin gene variants in acute (blue) and carrier (purple) isolates. Toxin variants were called based on previously reported alleles of the indicated toxins (Commons et al. 2014).

### *in vitro* biofilm formation is not correlated with carriage propensity

Bacterial biofilm formation is associated with numerous chronic infections and carriage of organisms on the mucosa. To determine whether there was a propensity for increased biofilm formation in the carrier or acute isolates, we tested the ability of 44 acute and 13 carrier isolates to form biofilms *in vitro* using a microtiter dish assay with a reference strain for comparison in each assay. Biofilm formation varied widely between isolates from very little biofilm formation to high biofilm formation, but there was no significant trend regarding differences in biofilm formation between carrier and acute isolates (Fig. S1).

### Genomic conservation in carrier and acute strains

Selected isolates (n=32, 18 acute samples, 11 carrier strains with an additional 2 sequenced “return” isolates) were submitted for both short (Illumina) and long (PacBio) read whole genome sequencing. Long read sequencing in conjunction with more stringent Illumina sequencing allowed for full annotation of the genomes and comparisons of both SNPs and indels as well as genome structure. Figure 3 is a circular plot showing comparative genomics for sequenced isolates including *emm* type concordant reference genomes. ∼60% of the genes in the pan-genome were considered the “core” (found in all 34 tested isolates although some had truncations) within this isolate cohort with the remaining ∼40% considered accessory gene content. The accessory genome content among the strains is high with both inter- and intra-*emm* type variation among the isolates. All rearrangements, duplications, or large indels associated with carrier isolates were also found in acute isolates.

**Figure 3.**
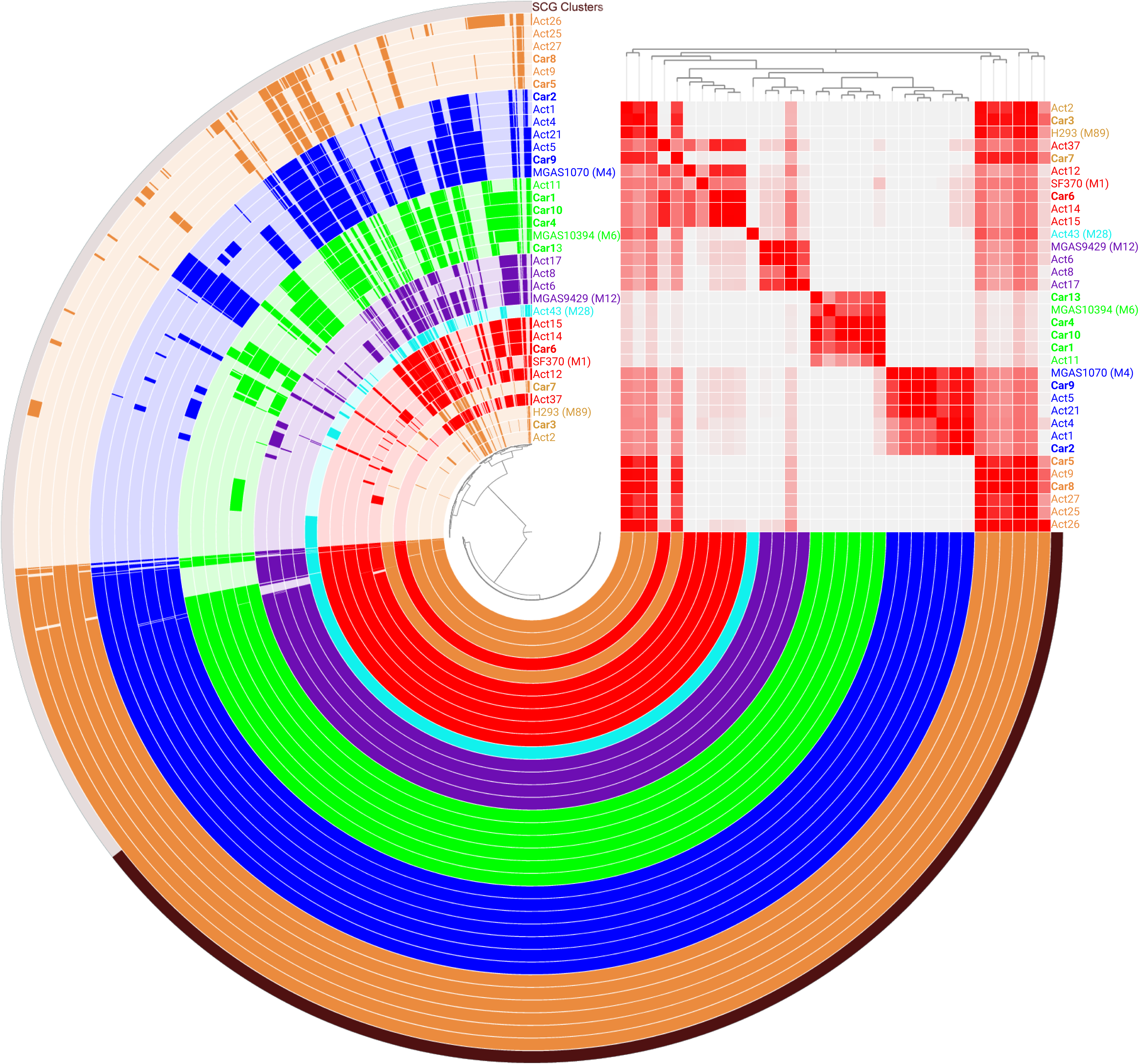
Anvi’o plot of sequenced isolates. Assembled genomes for each GAS isolate are shown alongside reference genomes for emm89 (brown) HG316453.2 (H293), emm1 (red) NC_002737.2 (SF370), emm28 (teal), emm12 (purple) NC_008021.1 (MGAS9429), emm6 (green) NC_006086.1 (MGAS10394), and emm4 (blue) NC_008024.1 (MGAS10750). ORFs, average nucleotide identity, and the inferred phylogeny are shown.

While no large genomic differences were immediately apparent to differentiate between the acute and carrier strains, this does not preclude SNPs, smaller indels, and virulence gene alleles from differentiating carriage and acute isolates. GAS contains a multitude of important virulence factors including toxins and capsule and several gene alleles previously identified to be associated with carriage in animal studies (21–23, 26). We examined nucleotide and amino acid sequences for some of these factors in our sequenced samples to determine whether particular allele variants or SNPs were associated with carriage or acute isolates. All strains contained between 2-5 toxin genes and no SNPs or indels were observed in these genes differentiating isolates from others of that same *emm type*.

Toxin allele variants of *smeZ, speA, speI, speC, speJ, speG, speH,* and *ssa* were determined based on previous literature (27). Our data did not reveal an obvious correlation between toxin gene presence and carriage or acute isolates (Fig. 2b).

The presence or absence of the selected virulence genes was almost completely *emm*-type dependent. Clumping factor A (*cfa*), for instance, was not found in any of the *emm* 6 isolates and *hasABC* genes were not found in any of the *emm* 4 or *emm* 89 isolates. Additionally, conservation of protein sequences generally matched in a serotype-specific manner rather than aligning by carriage versus acute designations. Amino acid polymorphisms in ScpA, for instance, were found in all isolates of the *emm* 89 and *emm* 6 isolates (Fig. 4).

**Figure 4.**
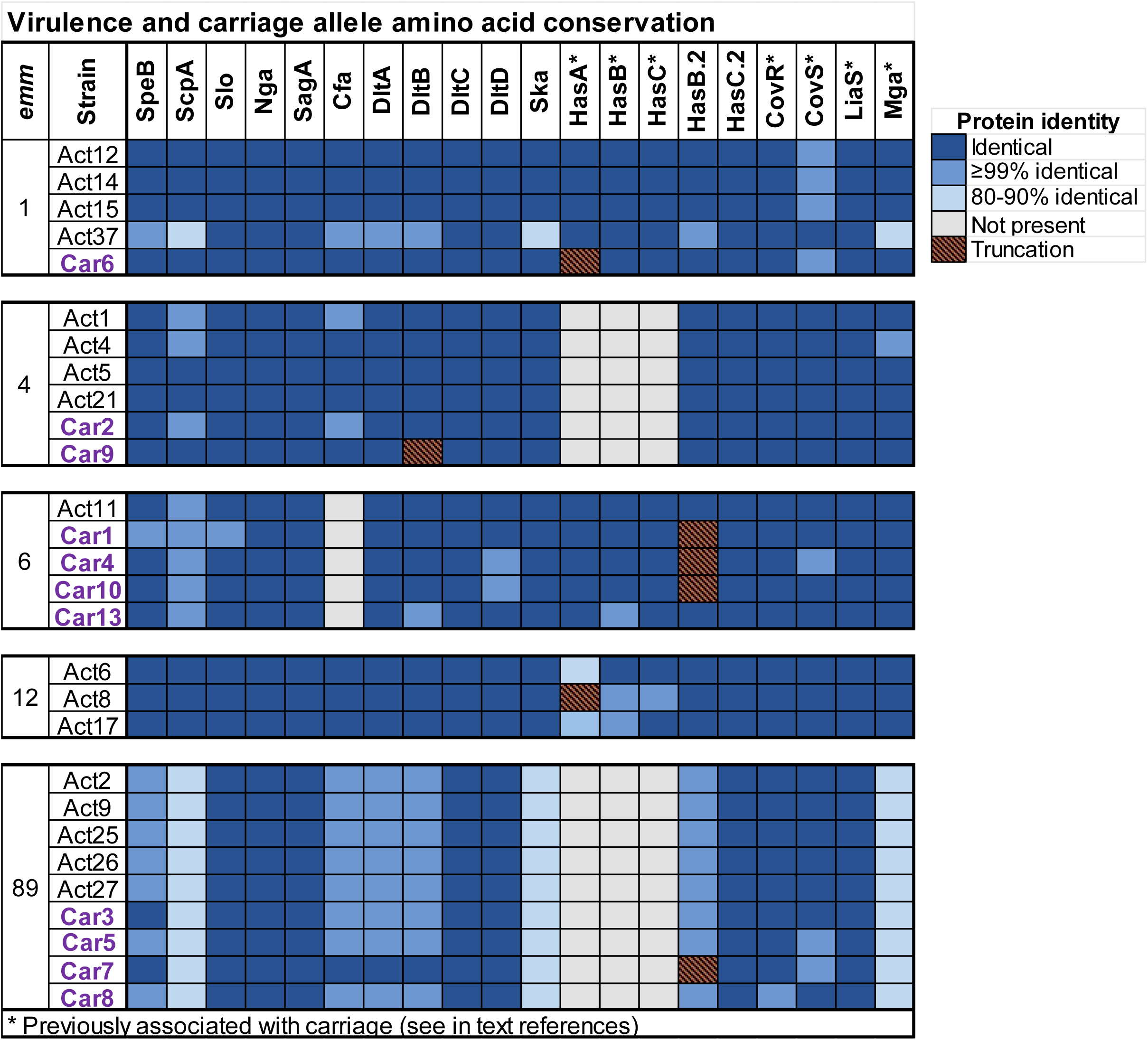
Changes in protein sequences are correlated to emm type over acute versus carrier profiles. Sequenced isolates are shown grouped by *emm* type and acute (black) versus carrier (purple). Protein sequences were compared to reference sequences of selected proteins. Blue color indicates percent amino acid identity to the closed allele. Gene truncations are shown in orange stripes. Gray indicates lack of coding sequence in that isolate.

SNPs in certain genes in GAS have been previously associated with carriage propensity using animal models of GAS colonization. Studies identified SNPs in a regulator, *liaS* (22), the capsule gene, *hasA* (26), and an adhesin protein *sclA* (23) primarily in carrier strains. When introduced into virulent strains of GAS, these mutations caused increased colonization of the murine nasal associated lymphoid tissue (NALT), as well as decreased phenotypes associated with virulence such as invasion into tissues and growth in human blood. None of the specific previously identified mutations were found in any of our carrier or acute isolates; although one carrier (Car13) did have a nucleotide insertion causing a frameshift in SclA (Table 3) and multiple carrier and acute strains were missing *hasA* altogether or had mutations or trunctions within it. A few amino acid polymorphisms (AAPs) were present only in carrier isolates from our study and not identified in any sequenced strains of GAS in the NCBI database (Table 3). Interestingly, one such polymorphism was found in two separate carrier isolates, Car4 and Car10 (both *emm* type 6) containing a SNP in the DltD gene resulting in an S364F change.

**Table 3:**
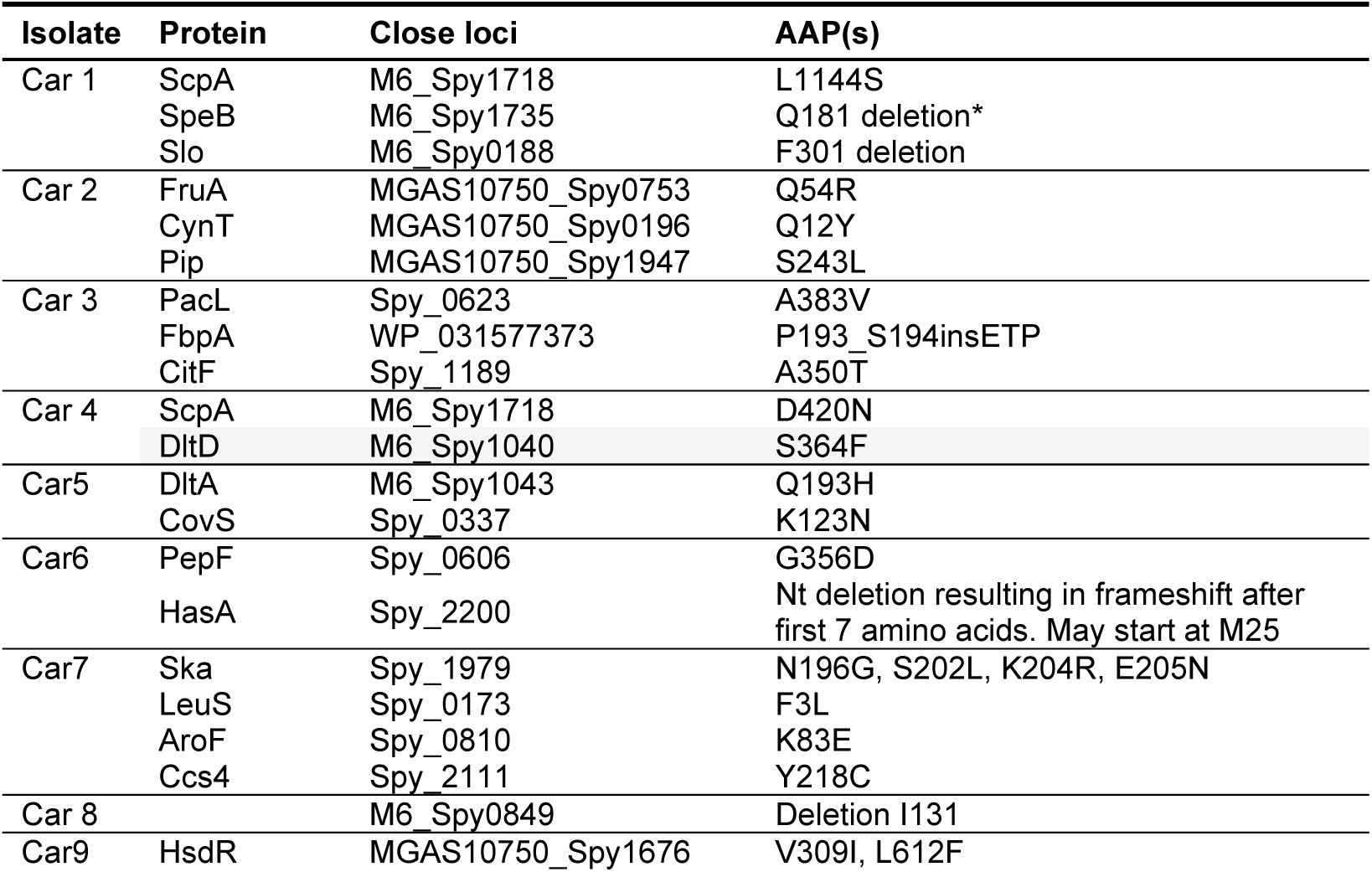
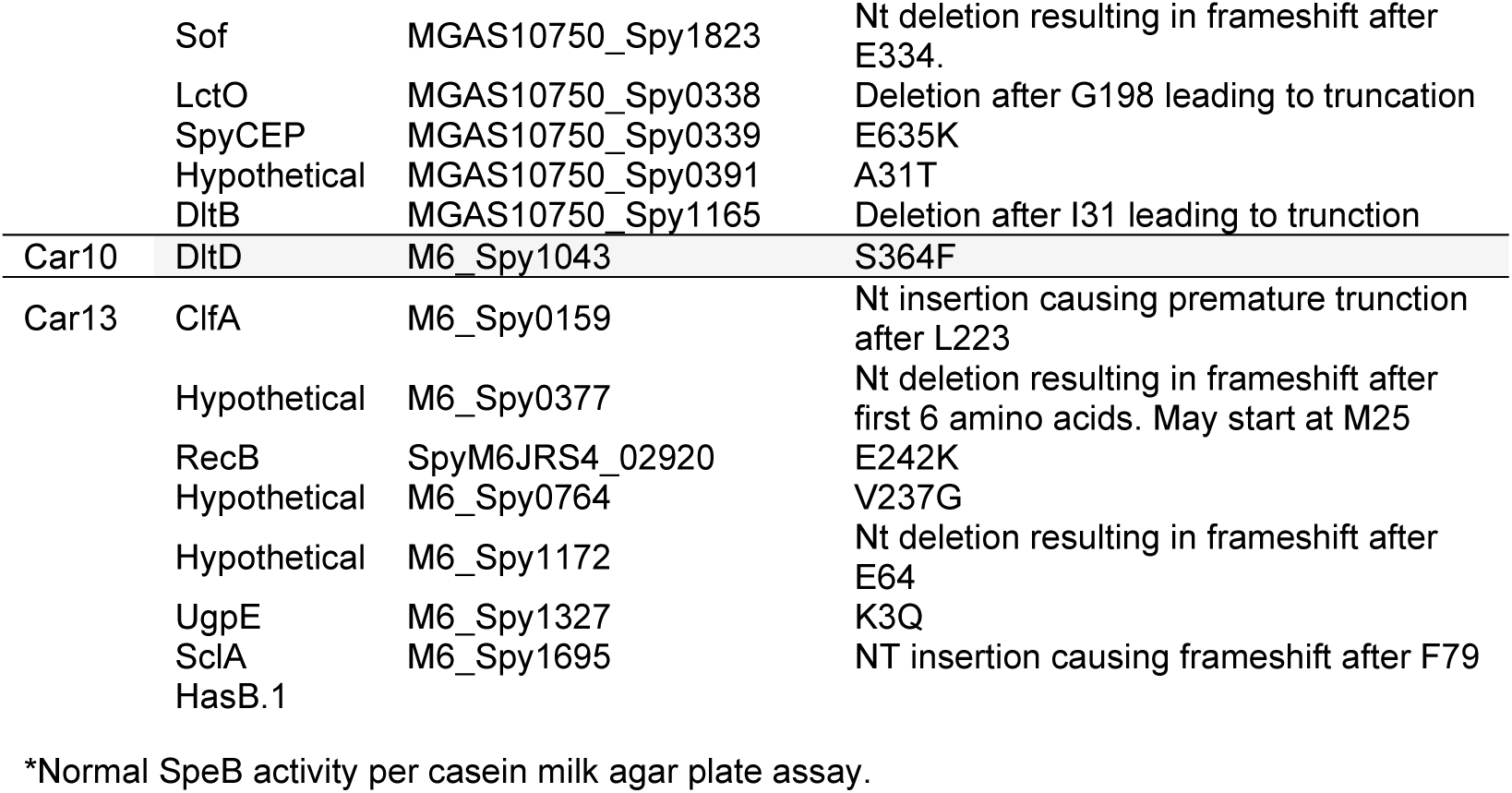
Carrier-only amino acid polymorphisms (AAPs)

Flores *et al*. previously identified a polymorphism found outside of an open reading frame in two carrier strains, a 12 bp deletion in the promoter of a major virulence regulator of GAS, *mga*. This change conferred a carrier-like phenotype when introduced into a typically virulent strain (21). The *mga* promoter region of all sequenced strains is shown in Figure S2. While some strains have deletions of up to 10 bp in this region compared to other strains, these deletions are present in both carrier and acute strains and were all *emm*-type dependent.

Besides comparing visit 1 acute and carrier isolates, we also compared genome sequences of two of the carriage isolates before and after antibiotic treatment. Car3-1 and Car 3-3 differed by two SNPs resulting in a Phe139Leu change in the gene *aapA (spy_1654)* and a T to G nucleotide at position 1,428,166 change in a non-protein coding region between the stop codons of adjacent genes. Additional differences were seen in assembly files such as duplications in the assembly but, when the genomic DNA was checked with Sanger sequencing in regions that did not match, the mismatches were shown to be the result of sequencing or assembly error. Corrected sequences for Car3-1 and Car3-3 are available as genbank files through NCBI’s GenBank database (reference numbers TBD).

Car13-1 and Car13-3 had two SNPs in rRNA genes; G to A at position 1741 and T to C at position 2969. Additionally, Car13-3 contained a nucleotide insertion of an additional A at position 1,271,084. This results in a trunction in HasB. Car13-3 does contain a complete HasB.2 and HasC.2 which have been shown to support capsule production in the absence of HasB (28). Additionally, Car13 contains a gene found in a few sequenced isolates of GAS annotated as a putative LPXTG adhesin homologous to M6_spy1173 (*smc*). In MGAS10394, this 1123 amino acid protein contains three full C terminal repeats of 91 amino acids. In Car13-1, this protein contains eight of these repeats and Car13-3 contains nine repeats (Figure S3), giving the resulting difference in genome size. The function of this protein is currently unknown.

### Transcriptional patterns of GAS strains during carriage and acute infection differentiate between disease states but are not predictive of future treatment failure and carriage

The lack of conserved carriage-specific SNPs or indels does not negate the hypothesis that carriage is driven primarily by polymorphisms that could change transcriptional patterns in carrier isolates. It is possible that SNPs and indels outside of open reading frames greatly affect the expression of key genes involved in immune evasion, antibiotic recalcitrance, and long-term carriage. Using patient swabs, we compared gene expression to determine whether “acute-specific” and “carriage-specific” transcriptional programs could be identified both before and after antibiotic treatment. We performed RNA-sequencing on swabs collected directly from patients during an acute infection. These swabs from the initial visit were split into groups based on whether the patient would go on to clear GAS following antibiotics (acute) or continue carrying GAS (treatment failure, carrier-1). Additional swabs were collected on a third visit from patients with treatment failure after antibiotics (carrier-3). Cultured liquid samples of many of the isolates were also run as comparators. In total, transcriptional profiling was done on 28 acute swabs, 9 carrier swabs (5 during the acute phase and 4 from the third visits), and 21 liquid cultures. This is the first description of GAS gene expression during an acute pharyngitis episode in human subjects.

To perform transcriptomics on disparate GAS isolates, we first created a fasta database file containing all coding sequences (CDS) from eight GAS reference genomes. 6 emm types (1, 2, 4, 6, 12, 28, 89) were represented, making up >85% of the emm type diversity found in our cohort. The final open reading frame (ORF) count in our pangenome was 2,326, making it approximately 35% larger than the average single GAS genome (1717 genes (29)). The number of differentially expressed genes (*q* ≤ 0.05, fold change ≥ 2) in each comparison is shown in Table 4 and Fig. 5. Gene expression from acute swabs were compared with isolates grown in liquid culture to delineate gene regulation during an acute pharyngitis episode (Fig. 5A, Table S1). As expected, a large portion of the genome was differentially regulated in this comparison. Many genes upregulated *in vivo* were involved in metabolic pathways such as histidine metabolism (Fig. 6), similar to what was observed in a murine model of necrotizing fasciitis (30). Fatty acid metabolism, in contrast, was downregulated in the swabs compared to growth in laboratory medium (Fig. 6). A similar decrease in fatty acid metabolism was observed for Group B Streptococcus during colonization of the vaginal tract compared to liquid culture (31).

**Figure 5:**
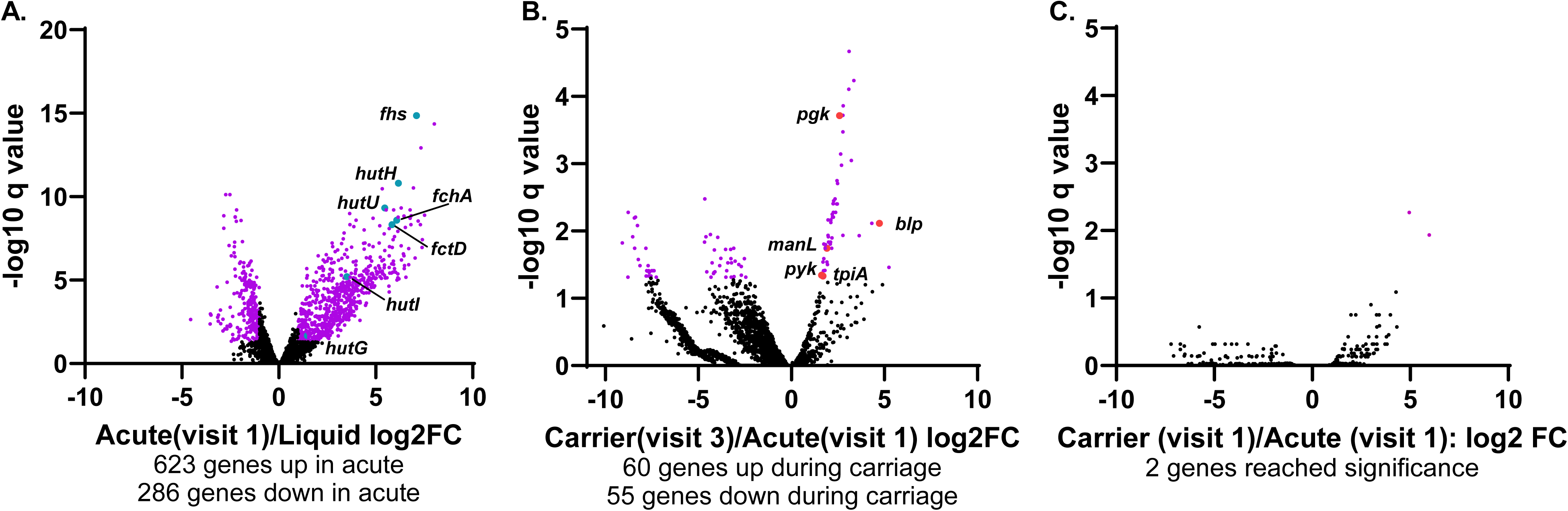
Significant changes in gene expression are seen between throat swabs and liquid samples and when comparing GAS during an active infection versus following a course of antibiotics. Throat swabs taken during an acute infection that resolved with antibiotics (acute, visit 1) were compared to (A) growth of isolates in liquid culture (liquid), (B) swabs taken during the asymptomatic phase (carrier, visit 3), and (C) swabs from an acute infection that would not go on to be resolved by antibiotics (carrier, visit 1). Genes in purple had q <0.05 and FC > Ι 2 Ι. Histidine metabolism genes are highlighted in teal. Alternative sugar metabolism and bacteriocins are highlighted in red. Acute visit 1 n=28, Liquid n=21, Carrier visit 1 n=5, Carrier visit 3 n=4.

**Figure 6:**
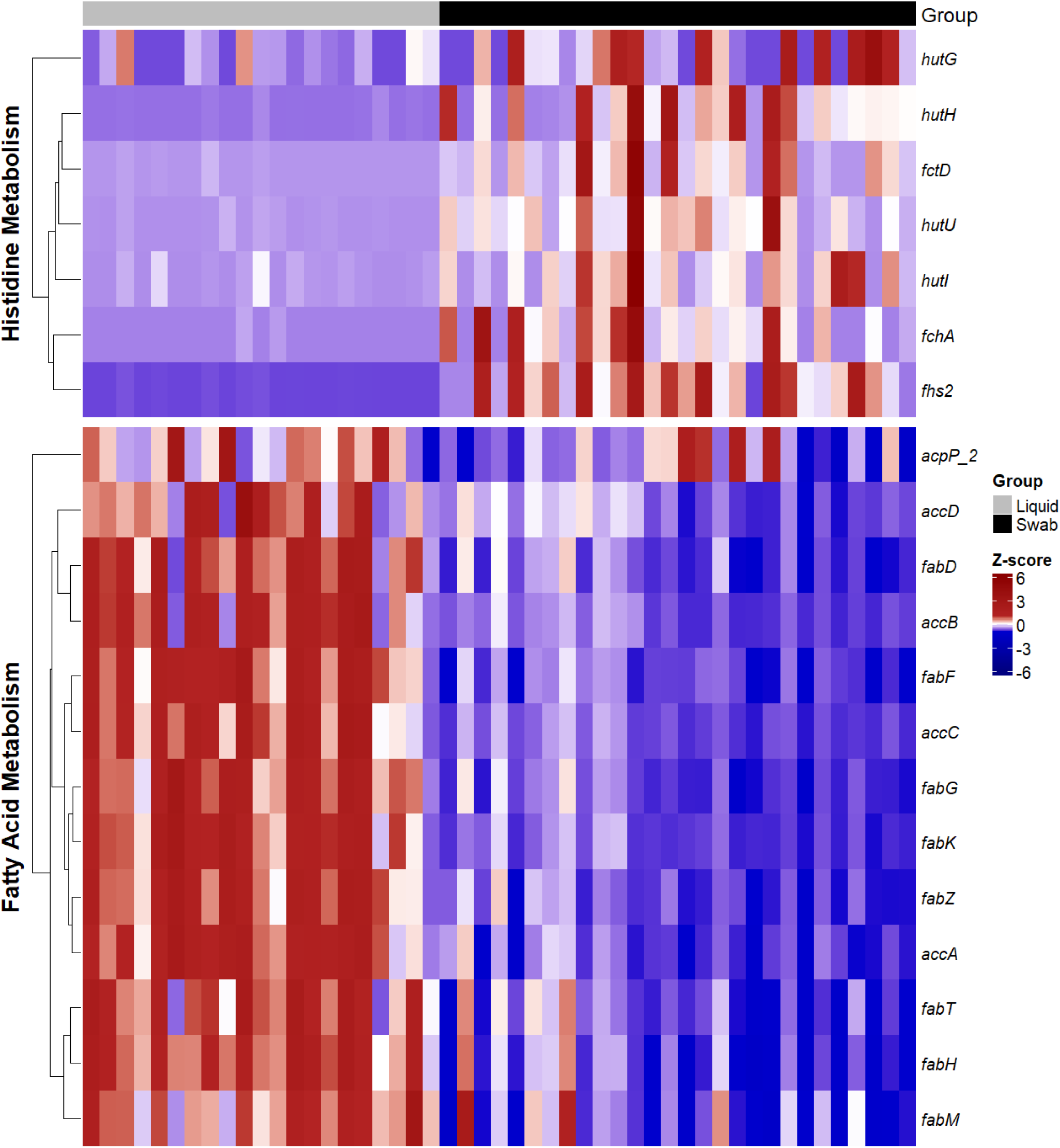
Heatmap of gene expression profiles in histidine and fatty acid metabolism genes. Row-scaled expression values for genes involved in Histidine Metabolism (top) and Fatty Acid Metabolism (bottom). Samples are divided into two groups, Liquid (gray, n=21) and Swab (acute, black, n-28). Gene expression values were standardized as Z-scores across each gene (row), such that values indicate the number of standard deviations from the gene’s mean expression. Genes within each metabolic group are clustered hierarchically, revealing expression patterns specific to sample types. The two metabolic pathways exhibit distinct expression signatures, highlighting differences in gene regulation between sample groups. The color gradient indicates relative expression levels from low (blue) to high (red). Data were processed and visualized using R (version 4.5.1) with the pheatmap package.

**Table 4:** Differentially expressed genes.

| Comparison* | Up | Down | Total |
| --- | --- | --- | --- |
| Acute1/Liquid | 623 | 286 | 909 |
| Carrier1/Acute1 | 2 | 0 | 2 |
| Carrier3/Acute1 | 60 | 55 | 115 |

\*Significant features at $Q < 0.05$ , $\log_2FC > 0$
Acute n = 28, liquid = 21, carrier 1 n = 5, carrier 2 n = 4

A comparison of GAS gene expression from swabs obtained during an acute episode (visit 1) and asymptomatic carriage (visit 3) showed differential expression of 115 genes, suggesting defined transcriptional programs associated with an active infection and carriage (Fig. 5B and Table S2). Glucose, the preferred sugar for GAS, is depleted in the nasopharynx. GAS has been shown to use alternative sugars including mannose, commonly found on glycans in the mucosa, in these cases (32). We observed a mannose transport gene (*manL*) and genes involved in glucose metabolism (*pgk, pyk, tpi*) upregulated during carriage (Fig. 5B). Recent data link alternative sugar metabolism to production of bacteriocins that lead to competitive interactions within the microbiota (32). During carriage, GAS must persist on the mucosal surface and interact with the host microbiota, leading to cooperative and competitive interactions. We also observed up-regulation of a class II bacteriocin (*blp*) during carriage indicating potential complex interactions between GAS and the nasopharyngeal microbiota (Fig. 5B).

Interestingly, many factors associated with virulence and inflammation such as enterotoxins, were not found to be significantly up-regulated during an active infection when compared to the carriage state in our swabs. Previous work using a long-term model of GAS carriage or sub-clinical infection found that genes such as *speB (spy1735), smeZ (spy1702), and scpA (spy1715)* were significantly upregulated during the first acute phase of infection (24). In our data set, no difference was seen in the expression of these genes comparing the acute and carrier state swabs. Other genes identified in that dataset as being correlated with acute infection such as *covS* and *sic* were up-regulated in our data set but not significantly. Many of the genes previously found in that data set to be positively correlated with carriage versus acute infection were conversely found to be higher in our acute dataset including *lplB* (*spy0758*) and higher although not significantly, regulator *ciaH* (*spy0947*), streptokinase (*ska, spy1684*), and *epf* (*spy0561*).

In contrast, GAS gene expression from swabs taken during an active infection (visit 1) which would later either be cleared (acute) or would remain following treatment (carriage) were generally indistinguishable (Fig. 5C, Table S3). When comparing these two groups, only two genes were significantly differentially expressed and, while they pass the threshold for statistical significance, the expression differences were driven by one or two swabs having very high expression levels in carriers, making them unlikely to be predictive of carriage. This indicates that GAS genomics and gene expression may not be the primary forces driving GAS transition to a carrier state, at least during an active infection. A lack of significant genomic or transcriptional changes in carrier and acute GAS isolates during the active pharyngitis episode as well as the epidemiology of chronic carriage in some children and lack of infection or carriage in others, indicate that external factors are likely to play a significant role in GAS infection and carriage.

## Discussion

We initially set out to determine whether conserved genomic or transcriptional changes in isolates of GAS could predict eventual treatment failure and carriage. To collect carrier samples, we used a longitudinal method that defined carriage as treatment failure following an acute pharyngitis episode but with resolution of inflammation. This necessarily provides a different set of samples than swabbing asymptomatic individuals, and has been used by several studies to define true GAS carriers (15–17).

One important characteristic for carriage is a throat culture persistently positive for GAS in a child who is asymptomatic (33), as is seen in our patient cohort. Using these isolates and material directly from the throat swabs, we present both whole genomic sequencing and transcriptomic analysis of GAS directly from patient samples for the first time. Previous studies have identified certain *emm* types as being more highly associated with acute infection and pharyngitis including M1, 2, 3, 4, 5, 6, 12, 28, 75, and 89 (34–36). In this small cohort, M6 and M89 were more predominant among carrier isolates. Our acute isolates fell into primarily 8 *emm* types: 1, 2, 3, 4, 6, 12, 28, and 89, in alignment with published data.

Previously published studies examining human samples collected over a period of one month have provided important preliminary data about oropharyngeal carriage of GAS (37, 38). Using cultured strains obtained from a surveillance study of Air Force cadets acutely and asymptomatically colonized with GAS (39), Beres *et al.* undertook whole genome sequencing (WGS) on carrier strains of GAS and compared them to strains causing acute infections (40). They identified that the core gene and prophage content of carrier strains was the same as a reference strain isolated from a patient with invasive disease; however, they hypothesized that allelic variation could contribute to differences between strains that cause acute infections versus those that are isolated from carriers. This study showed that isolates within the invasive or carrier groups differed by approximately 50-52 core single nucleotide polymorphisms (SNPs), and between groups, they differed by an average of 49 SNPs, indicating that the groups were not particularly differentiated. They also did not identify specific SNPs that differentiated all carrier strains from acute isolates.

Flores *et al*. determined that one polymorphism found in two of four carrier strains, a 12 bp deletion in the promoter of a major virulence regulator of GAS, *mga*, could confer a carrier-like phenotype when introduced into a typically virulent strain (21). Similarly, SNPs in genes such as a regulator, *liaS* (22, 41), the capsule gene *hasA* (26), and an adhesin protein *sclA* (23), when introduced into a virulent strain of GAS, caused increased colonization of the murine nasal associated lymphoid tissue (NALT) and decreased phenotypes associated with virulence such as invasion into tissues and growth in human blood. These data led to the hypothesis that genetic changes arising in key virulence factors during acute infection could be a major contributor to the carrier state and that mutations in virulent strains could give rise to carrier isolates presumed to have decreased pathogenic potential.

We used both long and short read genome sequencing to examine 18 isolates from acutely infected individuals, 11 isolates acquired from patients during an acute infection who remained carriers following antibiotic treatment, and 2 isolates from the same patients following antibiotic treatment. We examined whether particular amino acid polymorphisms (AAPs) were found at a higher frequency in carrier isolates. Searching specific genes previously proposed to be involved in carriage or virulence, we detailed AAPs found only in carrier isolates. Each of the 11 carrier isolates had AAPs found only in those isolates and not in strains causing acute infections in our study or sequenced strains in the NCBI database (Table 3). Two isolates (Car8 and Car10) had only a single specific AAP in the genes we examined, seven isolates had between 2 and 4 AAPs, and the remaining two isolates had 6-7 AAPs.

We have not ruled out the possibility that these AAPs contribute to carriage and some of them are found in known virulence factors ScpA, CovS, SpeB, Slo, and DltD.

Two genes had AAPs found in two different carriers. AAPs in ScpA were found in both Carrier 1 (L1144S) and Carrier 4 (D420N) and AAPs in DltD were found in both Carrier 4 (S364F) and Carrier 10 (S364F). Additionally, Car5 and Car9 had AAPs or trunctions in DltA and DltB respectively. The Dlt operon aids in incorporating D-alanine esters into teichoic acids and modulates bacterial surface charge. Inactivation of genes in this operon could lead to changes in resistance to cationic antimicrobial peptides (AMPs). It is possible that these identified AAPs provided a fitness benefit for carriage, possibly in competitive interactions with the host or resident microbiota. Truncation of DltB is predicted to increase susceptibility to AMPs and neutrophil killing (42) and decrease virulence (43) which may serve as a trade-off for carriage in a state of lower inflammation. The potential for increased carriage potential caused by these AAPs should be examined in more depth in the future.

As described above, some SNPs, indels, and AAPs have been previously identified as being potentially carriage associated (21, 22, 26, 41). A deletion in the promoter region of *mga*, an important regulator of gene expression, was previously shown to affect gene expression and host/pathogen interactions in carrier and acute isolates of GAS (21). When this deletion was inserted into an invasive strain, virulence was greatly decreased. We similarly found polymorphisms in the promoter of *mga* but, rather than being associated with carriage, identical deletions were found on all isolates based on *emm* type. All M4, 6, and 89 isolates contained deletions in the promoter while M1 and 12 did not, regardless of long term carriage outcome. This does not rule out *mga* promoter polymorphisms as playing a role in carriage, however, this suggests that there may be other factors at play. Additionally, this difference could be the result of sampling, as a higher proportion of carrier isolates were M6 and M89. An AAP in the regulator LiaS has also been shown to promote carriage over invasion (22, 41) but no mutations were identified in LiaS in any of our carrier isolates.

An animal model of pharyngitis in cynomolgus macaques was previously used to conduct long-term infection over a duration of several weeks. In these studies, temporal changes in GAS gene expression were examined over the course of the infection. The acute phase of infection was defined as days 4-23 following intranasal inoculation when acute infection indictors like tonsilitis and pharyngitis score and C reactive protein levels were high. The asymptomatic or carriage phase was defined as days 23-58 when these infection markers were low but GAS was still present. Both GAS gene expression (24) and host responses (38), were evaluated. These data provide an important framework to examine genetic programs that govern GAS infection and carriage, but also raise some questions. In these experiments, the carrier state is described as the period following an acute GAS infection in which symptoms of active infection (tonsilitis and pharyngitis) have decreased while GAS colonization remains. However, this may represent a phase of resolution rather than a true carrier state. Definitions of the carrier state have included a lack of serologic response as well as treatment failure, neither of which were demonstrated in carriage studies using the macaque model. In humans, the carrier phase is characterized by low antibody titers. In the macaque model, antibody responses against GAS antigens were shown to remain high at day 29 (44), a time point considered in later studies to be part of the carrier phase (24). GAS remains a human-restricted pathogen; nevertheless, animal model systems have provided important data on the GAS infection state and the changing genetic programs that occur over the course of an infection.

We observed carriage transcriptional programs that differed from gene expression during an acute infection. This was expected and likely influenced by several factors including differences in the inflammatory environment of the throat as well as changes in the microbiota composition following antibiotic treatment, both of which likely affect GAS transcription. Comparing gene expression in our human model with genes correlated with acute infection or carriage in the macaque model reveals both overlapping and non-overlapping patterns of gene expression. The acute infection in macaques resulted in high levels of expression of genes such as *smeZ, cbf, fasC, vicR,* and *covS* and acute infection parameters were negatively correlated with *covR, ska, cfa,* and *hlyX* (24). In our study, *smeZ* was similarly upregulated ∼10 fold while *ska* was similarly downregulated during acute infection compared to liquid culture. *cbf*, the *fasBCA* system, *vicRK*, *cfa, hlyX*, and *covRS* expression, however, showed no change when comparing bulk liquid cultures from swab samples. However, these are not particularly equivolent comparisons as we are comparing liquid culture to throat swabs, where the macaque experiment was looking at expression levels compared to clinical parameters.

Virtanevna et al. also showed correlations between specific GAS genes and the asymptomatic infection stage in macaques (24). They found that several factors were down or up-regulated on day 32 (asymptomatic infection) versus day 16 (acute infection). Some of the most highly down-regulated genes observed in their study when comparing later infection stages to early include *hlyX* (spy0317), *atpF* (spy0577), *proB* (spy1372), and *accA*, (spy1485), among others. While these genes were also all down-regulated in our asymptomatic carrier conditions, only one of them (*accA*) reached significance. Some of the most highly up-regulated genes in the day 32 samples compared to day 16 included *adrC* (spy 0078), *fusA* (spy0232), *tcyA* (spy0982), and *hup* (spy1223). *hup* and *fusA* were also significantly up-regulated in our data set during carriage but *adrC* and *tcyA* were not significantly differentially expressed.

Our study has several limiting factors. While we have a broad range of isolates more representative of the diversity of strains in the population, it also makes transcriptomic differential expression comparison more difficult as the strains being compared are not genetically identical like in the macaque study. This creates the need for a pan-genome to examine overall transcriptional profiles.

Because some genes are found in particular *emm* types more than others and there is an over-representation of *emm* type 6 in our carriage isolates, this leads to skewed statistics showing possible overexpression of *emm-*type specific genes during carriage. We have focused much of the discussion on genes found in the core genome in most of the isolates, but care should be taken when interpreting the expression patterns of *emm-*type specific genes in the dataset.

Additionally, patient cohorts were made up of children who all experienced an acute pharyngitis episode, not children without a history of oropharyngeal symptoms. In our protocol, it may be difficult to detangle the role of antibiotic pressure on bacterial mutations or gene expression. We expect that antibiotic treatment would strongly affect gene expression of the bacteria in the short term, but this pressure would most likely be negligible weeks after antibiotics were completed. Antibiotic usage will also influence the microbial communities present and changes in the local oropharyngeal microbiota likely influence GAS transcriptional patterns as well.

Despite these limitations, our study is the first to report GAS transcriptomic data from a human cohort using oropharyngeal swabs. We can define transcriptional profiles associated with acute infection when compared to growth in laboratory medium (Fig. 5A and Table S1) or compared with swabs taken post-antibiotics (Fig. 5B and Table S2). However, we did not observe any major differences between swabs taken during an acute pharyngitis episode comparing those that would later have treatment failure versus those in which infection would be resolved (Fig. 5C and Table S3). While this study does not determine whether GAS genomics or transcriptomics primarily drive carriage or whether host or native microbiota interactions play a bigger role, it does give important insight into these processes from a human infection perspective.

## Materials and Methods

### Swab collection

Children were recruited from four pediatric clinics in Madison, WI, USA and eligible for this study if they were between 5 and 15 years of age, had a diagnosis of acute pharyngitis, and had not been treated with antibiotics in the previous 30 days. Children were excluded from this study if they had an allergy to β-lactam antimicrobials or were treated with an antibiotic other than a β-lactam. A swab was obtained from the posterior pharynx (tonsillar pillars and posterior pharyngeal wall) of the patient. A rapid antigen detection test (RADT) was used for initial diagnosis. If the RADT was positive, the child and family were informed by the research staff that the child was eligible for the study and if the family and patient agreed to participate, signed informed consent and assent were obtained. At that point, two additional swabs were taken. One was sent to the laboratory to be cultured for GAS using standard methods. The cultured isolate was preserved. The additional swab was stored immediately at -80 °C. Treatment was undertaken with an appropriate dose of either amoxicillin or penicillin V for 10 days at the discretion of the provider. The patient then returned for follow-up 13-14 days after initiation of treatment. If the child was asymptomatic and lacked apparent inflammation of the pharynx or tonsils, a double swab was used to sample the posterior pharynx. One was sent to the laboratory to be cultured for GAS using standard methods. The cultured isolate was preserved. The additional swab was stored immediately at -80 °C. If the follow-up swab was negative for GAS on culture, the patient was deemed to have presented with an acute infection. If the follow-up swab was positive, the second swab was assigned as probably carrier. An additional set of swabs was then obtained 1 week later, i.e around 14 days after completing antibiotics. One was sent to the laboratory to be cultured for GAS using standard methods. The cultured isolate was preserved. The additional swab was stored immediately at -80 °C. If the swab was still positive, the subject was deemed a confirmed-carrier and the swab and cultured isolates were stored at -80°C. All cultured isolates from a given patient were tested by PCR to demonstrate the same *emm* type in the first and last swabs. Only those with concordant *emm* type were compared for differential gene expression. Swabs were collected under protocol #2015-1456-CP016 approved by the University of Wisconsin Institutional Review Board (IRB).

### *emm* typing

*emm* typing of isolates was done as previously published (25, 45). Genomic DNA from isolated cells was amplified using published primers *emm* Sense: TATTGGCTTAGAAAATTAA and *emm* Antisense: GCAAGTTCTTCAGCTTGTTT (25, 45). PCR amplified DNA was sequenced and sequences aligned to the closest *emm* type using NCBI Blast. The nearest sequence match was analyzed for SNPs in the first 180 residues and *emm* types were identified if no SNPs were found.

### Biofilm formation

Biofilm assays were done as previously described (46, 47) with minor modifications. Bacterial cultures were grown overnight in Todd Hewitt medium + 2% yeast extract (THY) statically at 37°C. Cultures were diluted to an OD600 of 0.05 in fresh THY + 0.5% glucose and 1 mL of the cell suspension was added to wells of a 24 well plate in at least technical triplicate and incubated at 37°C. After 24 hours, the culture medium was removed, and the wells were washed 3 times with 1 mL phosphate buffered saline (PBS) to remove unattached cells. 1 mL of 0.1% crystal violet solution was added to each well and incubated at room temperature for 10 minutes. Excess crystal violet was removed, and the wells washed another 3 times with 1 mL PBS. Remaining stain was dissolved in 1 mL 95% ethanol and absorbance at OD590 was measured. The average of ≥2 technical replicates was counted as a single point and the experiment was performed for each strain in at least biological triplicate on different days. The lab strain NZ131 was included on each plate as an internal control for variability and values are shown relative to NZ131 on each plate. Values shown are the average and standard deviation of the biological replicates.

### Minimum inhibitory concentration tests

MIC assays were done using MIC Test Strips (MTS, Liofilchem) for clindamycin (0.016-256 µg/mL, 92072), amoxicillin (0.016-256 µg/mL, 92021), penicillin G (0.002-32 µg/mL, 92103), and erythromycin (0.016-256 µg/mL, 92051). Frozen culture stocks were streaked onto Blood Agar Plates (BAP, Hardy Diagnostic, A10). Individual β hemolytic colonies were grown overnight in THY. Per manufacturer’s instructions, overnight cultures were diluted to OD600 = 0.5 and sterile swabs were used to streak lawns of bacteria on BAPs. Antibiotic strips were added, and the plates were incubated for the manufacturers’ recommended time and temperature. MIC values were recorded per manufacturer’s instructions.

Resistance to antibiotics were defined as follows: penicillin MIC ≥4 mg/L, amoxycillin MIC ≥ 1 mg/L, clindamycin MIC ≥ 1 mg/L, erythromycin MIC ≥ 1 mg/L.

### Preparation of DNA for Illumina short-read whole genome sequencing (WGS)

Genomic DNA was prepared using overnight cultures of GAS grown in THY at 37°C statically.

Cultures were pelleted and pellets were resuspended in 150 µL lysis buffer (10 mM Tris, 1 mM EDTA, pH 8.0 with 1% w/v SDS) and incubated at 55°C for 15 min. 1/5 volume 3 M NaAc was added and the cells were incubated at 65°C for 30 min and then incubated on ice for 1 hour. Samples were centrifuged at 15,000 xg for 15 minutes. DNA in the supernatant was precipitated by adding 2 volumes of ice cold 100% ethanol followed by incubation at -80°C for at least 60 minutes. DNA was pelleted by centrifugation at 15,000 xg for 10 minutes. The supernatant was removed and the pellets were air dried. The resulting DNA pellet was resuspended in 100 µl water and the concentration and purity measured on a Nanodrop OneC (Thermo Fisher). DNA libraries for WGS were prepared from 500 ng of gDNA in-house using the Illumina Nextera Flex DNA library preparation kit (Illumina 20018704) per manufacturer’s instructions using the Nextera DNA CD Indexes (ThermoFisher 20018707). Final libraries were sent to the University of Illinois at Chicago Research Resources Center for quality control, sequencing, and analysis.

### Preparation of DNA for PacBio long-read whole genome sequencing

High molecular weight (HMW) DNA was prepared for Pac Bio sequencing libraries using the HMW DNA Extraction kit for Tissue and Bacteria (NEB T3060) according to manufacturer’s instructions for low input Gram positive bacterial samples with some modifications. Overnight cultures grown in THY were spun down to an equivalent of OD600 = 4.0. Pellets were resuspended in 150uL of STET buffer with freshly added lysozyme at 50 mg/mL. Homogenization was carried out according to kit manual, using the thermal mixer protocol at 1400 rpm. DNA was sheared using the Covaris g tubes (Covaris 520079) per manufacturer’s instructions using 100 µL/sample and spinning at 9400 rpm for 2 minutes on each side. Sheared DNA was then cleaned up using the 0.45X AMPure PB beads (PacBio 100-265-900) and Elution Buffer (PacBio 101-633-500) following the PacBio protocol. The quality and quantity of DNA was checked using a Nanodrop OneC and only samples with A260/280 ratios of >1.8 were used to make libraries. PacBio libraries were made in-house with the SMRTbell Express Template Prep Kit 2.0 (PacBio 100-938-900) using the SMRTbell Barcoded Adapter Plate 3.0 (PacBio 102-099-200) per manufacturer’s instructions. Libraries were sent to the University of Illinois at Chicago (UIC) Genomics Core facilities for quality control and sequencing on a Sequel II SMRT Cell.

### Genome assembly

Prior to assembly, PacBio data were filtered to remove reads shorter than 1000 bp and trimmed to remove adapters using Porechop v0.2.3 (48). Filtered and trimmed reads from each isolate were individually assembled using Flye v2.9 (49) with the PacBio HiFi option and command line options genome-size=2m and asm-coverage=100. Illumina reads sequenced from the same isolate were aligned to assembled contigs using the Burrows-Wheeler Aligner MEM algorithm (BWA MEM) v0.7.17 and the mpileup output from the alignment was processed to generate a polished assembly for each isolate (50). Each polished assembly was used for further alignment and polishing, repeating until no changes could be detected. Prokka (51) was used to detect open reading frames (ORFs) and perform functional annotation of the final polished assembly for each isolate.

### Pan-genome analysis

Assembled genomes and functional annotations for each GAS isolate were loaded into Anvìo (52) alongside reference genomes HG316453.2, NC_002737.2, NC_006086.1, NC_008021.1, and NC_008024.1 using the anvi-gen-contigs-database, anvi-import-functions, and anvi-gen-genomes-storage programs. The pan-genome analysis was run using the anvi-pan-genome program. Average nucleotide identity was computed using the anvi-compute-genome-similarity program with the pyANI option (53). Single-copy core genes (SCG) were extracted from the pan-genome analysis, concatenated and aligned using the anvi-get-sequences-for-gene-clusters program for subsequent multi-locus phylogenetic analysis. Amino-acid positions with gaps in >50% of genomes were excised from the multi-locus alignment using trimAl (54). FastTree (55) was used to infer the phylogenetic relationship from the multi-locus sequence alignment and was loaded into Anvìo using the anvi-import-misc-data program. A figure was generated using the anvi-display-pan program to display ORFs, ANI, and the inferred phylogeny, with colors added manually to each track to highlight the *emm* types for each genome.

### RNA extraction and depletion of eukaryotic and ribosomal rRNA

For liquid samples, overnight cultures of GAS strains were grown in THY medium. In the morning, cultures were diluted 1:10 in fresh THY medium and grown at 37 °C static until the OD600 reached 0.4-0.6. Cells were then spun down and resuspended in 500 µL DNA/RNA Shield (Zymo) before storage at -80 °C until extraction. When ready, samples were thawed at room temperature, centrifuged to pellet the cells and the DNA/RNA Shield removed. Pellets were then resuspended in 200 µL lysis solution (TRIzol Max Bacterial Enhancement Reagent (ThermoFisher 16122012) which had been pre-heated to 95 °C mixed with 1 mg/mL lysozyme and 50 U/mL mutanolysin) and moved to a MagMAX prime bead beating tube (ThermoFisher A58155). For swabs, 500 µL DNA/RNA Shield was added to frozen swabs and then vortexed for 30-60 seconds. The tubes containing the swabs were then sonicated in a sonicator bath for 5 minutes to help with removing adherent cells. Following sonication, swabs were again vortexed for 30-60 seconds and then the tubes were spun down for 5 minutes at 4 °C. The supernatant and swab were removed and the resulting pellet resuspended in 200 µL lysis solution and moved to MagMAX bead beating tubes as above.

All samples were placed in an OMNI Bead Ruptor 12 (Omni International, 19-050A) and beat twice for 4 minutes each with a 30 second break in between. The mixtures were then incubated at 37 °C for 30 minutes to allow additional enzymatic lysis and then incubated at 95 °C for 5 minutes. RNA was then collected using the TRIzol Max Bacterial RNA Isolation kit (ThermoFisher 16096-040) per manufacturer’s instructions including the phase separation and RNA precipitation steps. RNA pellets were air dried and resuspended in 45 µL DEPC-treated water. The Turbo DNA-free kit (ThermoFisher AM1907) was used per manufacturer’s instructions to remove any remaining DNA in the sample. The final RNA quantity and quality were measured on a NanoDrop OneC.

Depletion of eurkaryotic RNA was done on the RNA from the swab samples using the MICROEnrich kit (ThermoFisher AM1901) per manufacturer’s instructions. The final precipitation step was skipped and the samples were taken immediately into the MICROBExpress kit protocol (ThermoFisher AM1905) per manufacturer’s instructions to remove bacterial rRNA. For liquid cultures, bacterial rRNA was removed using the same MICROBExpress kit using 2-10 µg starting RNA in ≤15 µL total volume.

### RNA-sequencing library preparation and sequencing

RNA-seq libraries were prepared in house using two different protocols. Libraries from swabs processed before 2020 were prepared using the Kapa Stranded RNASeq Library Preparation kit (KK8400) per manufacturers instructions for purified mRNA with input range of 10-100 ng total. RNA was fragmented at 94°C for 6 minutes with random primers and separate illumina adapters were ligated onto each samples. The libraries were then frozen at -20 °C and sent to the University of Illinois at Chicago Genomics facility for Tapestation QC and Quibit quantification prior to sequencing on an HiSeq 2500. Libraries from swabs processed after 2020 were made using the the NEB Next Ultra II Directional RNA library prep kit (NEB E7765) per the manufacturer’s protocol for purified mRNA. mRNA input amounts ranged from 1-100 ng total. Briefly, rRNA depleted RNA in was first fragmented at 94°C for 10 minutes with random primers. First and then second strand cDNA was synthesized, followed by cDNA purification using SPRIselect beads. Ends were prepped and NEBNext adaptors were ligated to cDNA strands. Samples were again purified using SPRIselect beads, followed by PCR enrichment of adaptor ligated DNA using unique indexed forward and reverse primers. Samples were purified once more using SPRIselect beads, and library quality was analyzed using a NanoDrop (Thermo). The libraries were then frozen at -20 °C and sent to the University of Illinois at Chicago Genomics facility for Tapestation QC and Quibit quantification prior to sequencing on an Illumina NovaSeq X 10B, 2x150bp reads.

### RNA-seq quantification and differential analysis

A custom FASTA database was generated containing all of the coding sequences (CDS) from reference genome GAS ATCC 12344 and CDS from HG316453.2, NC_002737.2, NC_006086.1, NC_007296.2, NC_008021.1, NC_008022.1, and NC_008024.1 that could not be found in the ATCC strain using BWA MEM v0.7.17 (50). Raw RNA-seq reads were aligned to this custom database using BWA MEM. The expression level of gene features (CDS regions from the reference) were quantitated using FeatureCounts v2.0.3 (56) as raw read counts of the stranded libraries. Differential analysis of quantitated gene features as compared with treatment (acute vs carrier) or sampling method (liquid vs swab) was performed using the software package edgeR (57, 58) on raw sequence counts. Prior to analysis, the data were filtered to remove any features that had less than 5 total counts summed across all samples. Data were normalized as counts per million and an additional normalization factor was computed using the trimmed mean of M-values (TMM) algorithm. Statistical tests were performed using the “exactTest” function in edgeR. Adjusted p-values (q-values) were calculated using the Benjamini-Hochberg false discovery rate (FDR) correction. Significant gene features were determined based on an FDR threshold of 0.05 (5%).

### Statistical analysis

Statistical analyses are described in the figure legends or in more detail in the methods. Fisher’s exact test was used to determine statistical significance for *emm* distributions. Students t test was used to determine differences between carrier and acute biofilm formation. ** p value < 0.01.

## Supplemental material

Supplemental figures 1-3 show *in vitro* biofilm formation of isolates, *mga* promoter alignment, and protein domain architecture of Smc (M6_Spy1173) from sequenced MGAS10394 and Car13-1 and 13-3 for comparison respectively. Supplemental tables 1-3 are tables showing differentially expressed genes from RNA sequencing data presented in Figure 5.

## Acknowledgments

This work is supported by the National Institutes of Health (1R21AI147502 Cook/Wald MPI). Bioinformatics analysis in the project described was performed by the UIC Research Informatics Core, supported in part by NCATS through Grant UL1TR002003. We acknowledge undergraduate students in the Cook lab for performing biofilm assays shown in the supplemental information: Mio Ito, Lindsay Thomas, Jenna Battaglia, Nicole Remes, Courtney Fu, Diana Maneri, Mariia Samofalova, Isabella Caridi, James deBlasi, Teresa Scotto, Laura Bousaid, Sarah Patach, Keara O’Donnell. All authors approved the manuscript prior to submission.

---

**Figure S1.**
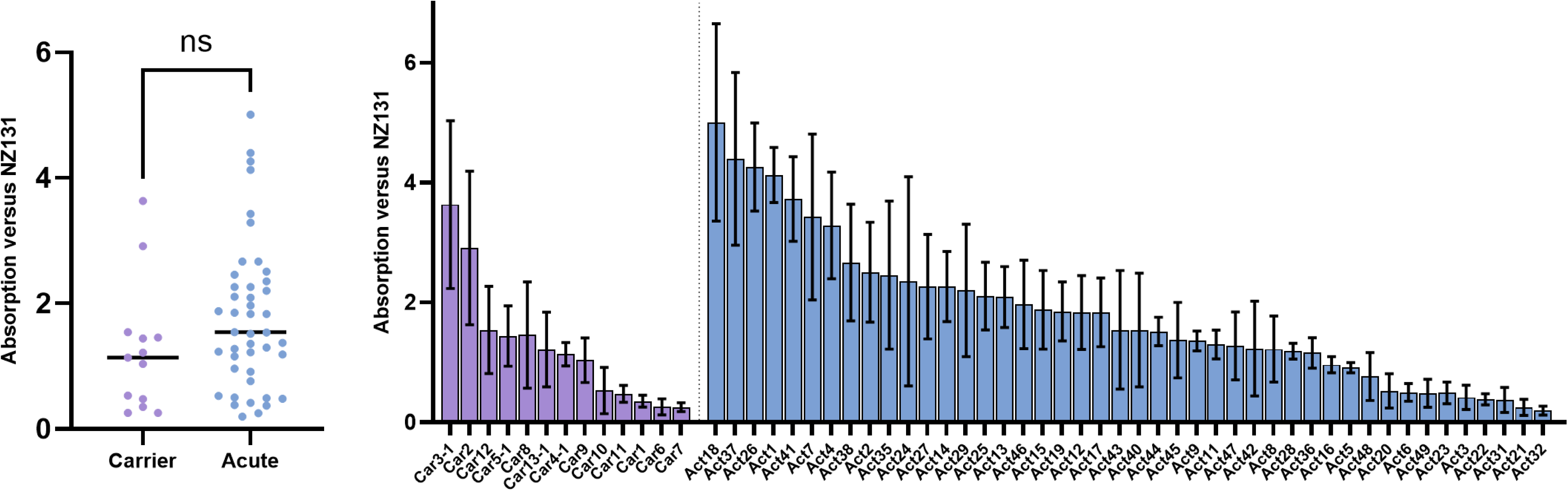
***in vitro* biofilm formation is not associated with carriage propensity.** Biofilm formation on abiotic surfaces was measured using crystal violet staining. Crystal violet absorption is given as a ratio to GAS strain NZ131 which was used as an internal control for each assay. The absorbance by group is shown (**a**) as are values for all individual isolates (**b**). At least three biological replicates and two technical replicates were used for each data point. The average and standard deviation are shown. Students *t* test used to analyze data. ns = not significant.

**Figure S2.**
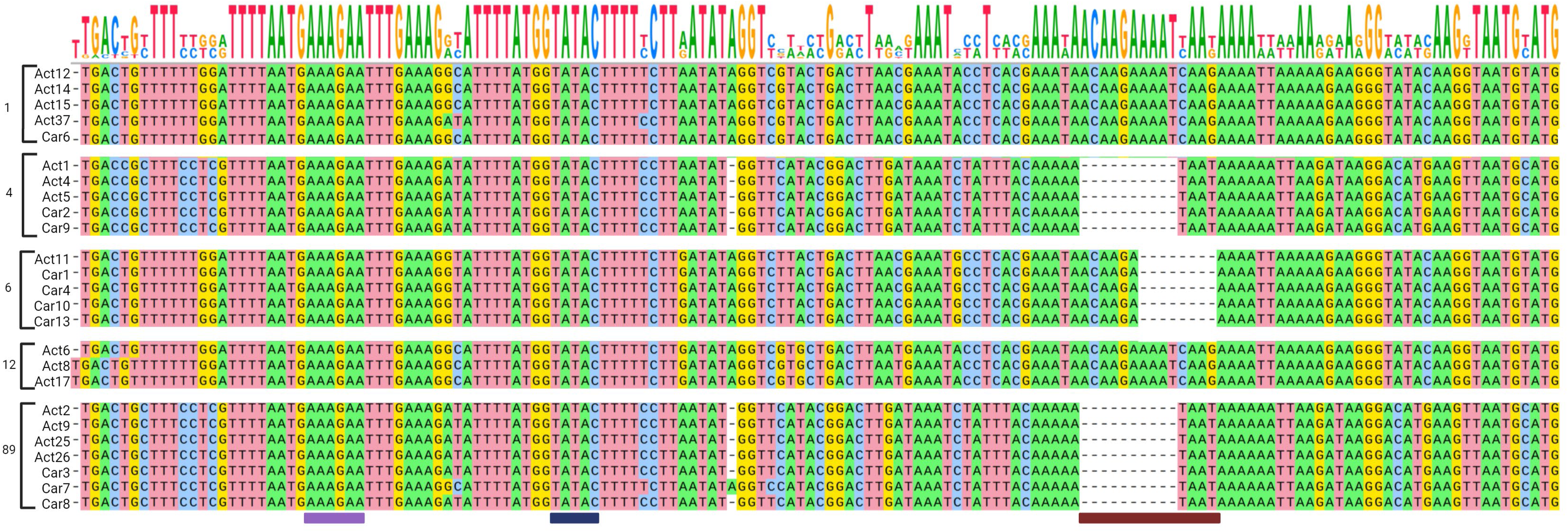
Alignment of the *mga* promoter region in sequenced strains. Purple and dark blue boxes represent the -35 and -10 sites respectively. Maroon bar indicates regions of deletions observed, which are conserved based on emm type in these sequenced isolates.

**Figure S3.**
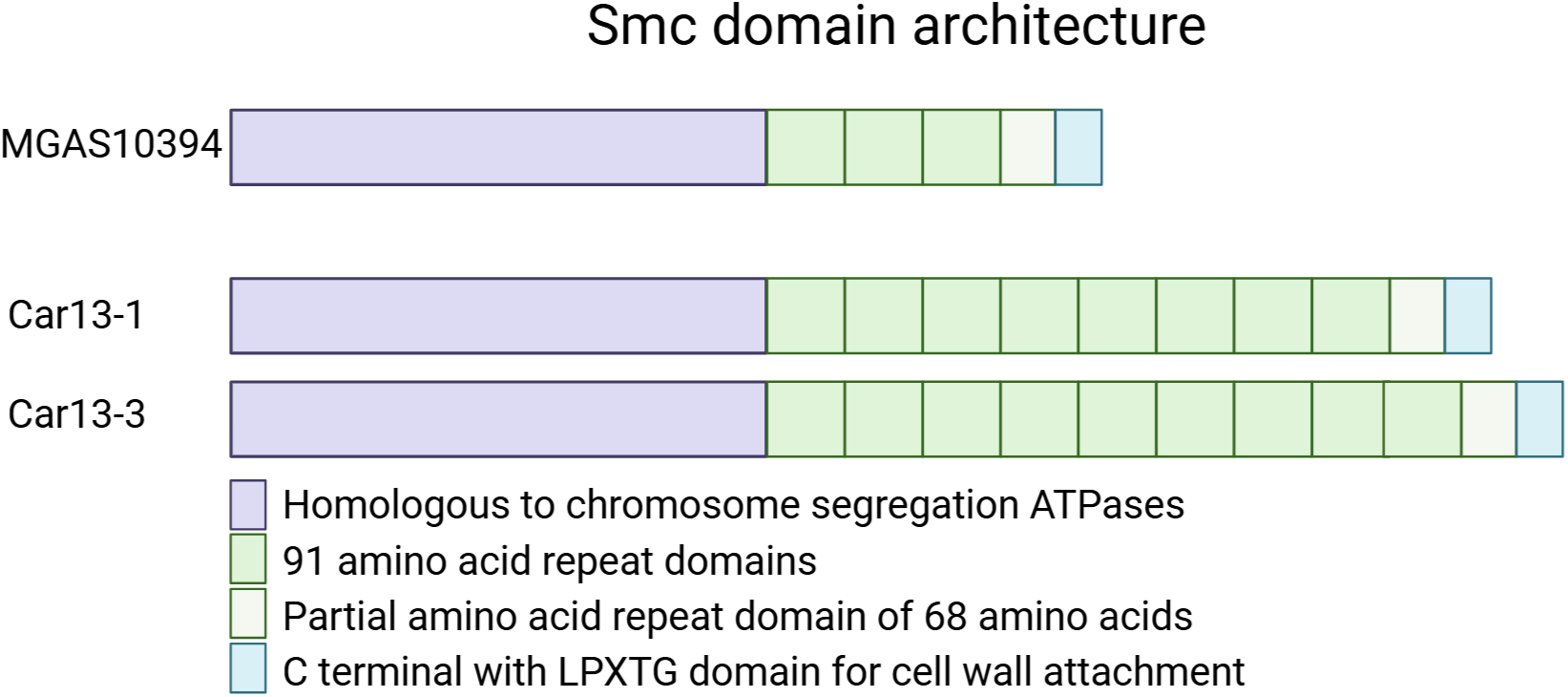
**Domain architecture of Smc**. The closest homolog to Smc is M6_Spy1173 from MGAS10394. Smc is found in Car13-1 and Car13-3 but the number of repeat domains differs between the three strains as shown. The purple region of the protein is predicted homologous to a chromosome segregation ATPase while the green and blue sections are predicted to be surface exposed adhesins with an associated LPXTG domain in the blue region.

